# MNPmApp: An image analysis tool to quantify mononuclear phagocyte distribution in mucosal tissues^a, b^

**DOI:** 10.1101/2021.09.27.461889

**Authors:** Catherine Potts, Julia Schearer, Dominic Bair, Becky Ayler, Jordan Love, Jennifer Dankoff, Paul R. Harris, Dominique Zosso, Diane Bimczok

## Abstract

Mononuclear phagocytes (MNPs) such as dendritic cells and macrophages perform key sentinel functions in mucosal tissues and are responsible for inducing and maintaining adaptive immune responses to mucosal pathogens. Positioning of MNPs at the mucosal epithelial interface facilitates their access to luminally-derived antigens and may regulate MNP function through soluble mediators or surface receptor interactions. Therefore, accurately quantifying the distribution of MNPs within mucosal tissues as well as their spatial relationship with other cells is important to infer functional cellular interactions in health and disease. In this study, we developed and validated a MATLAB-based tissue cytometry platform, termed “MNP mapping application” (MNPmApp), that performs high throughput analyses of MNP density and distribution in the gastrointestinal mucosa based on digital multicolor fluorescence microscopy images and that integrates a Monte Carlo modeling feature to assess randomness of MNP distribution. MNPmApp identified MNPs in tissue sections of the human gastric mucosa with a specificity of 98.3 ± 1.6% and a sensitivity of 76.4 ± 15.1%. Monte Carlo modeling revealed that mean MNP-MNP distances were significantly lower than anticipated based on random cell placement, whereas MNP-epithelial distances did not significantly differ from those of randomly placed cells. Interestingly, *H. pylori* infection had no significant impact on MNP density or distribution with regards to MNP-epithelial distances or MNP-MNP distances in gastric tissue. Overall, our analysis demonstrates that MNPmApp is a useful tool for unbiased quantitation of MNPs and their distribution at mucosal sites.

## Introduction

Immune cell interactions with their tissue environment can shape immune function and responses to infection and injury. Mononuclear phagocytes (MNPs) consist of blood monocytes and tissue-resident dendritic cells (DCs) and macrophages that play key roles as sentinel and antigen presenting cells (1). In previous studies on MNPs in the gastrointestinal mucosa, we and others have demonstrated that MNPs respond to environmental cues from both mucosal epithelial cells and stromal cells and that microenvironmental conditioning defines MNP function (2–6). Quantifying local cellular interactions between MNPs and other cells within the tissues is important to fully understand these functional immune networks in health and disease.

Microscopic images of tissues provide crucial information on immune cell density and distribution. Automated quantitative analysis of immunofluorescently labeled histological images, also termed tissue cytometry, enables unbiased high-throughput processing of digital imaging data (7). Multiple image analysis software packages from commercial and non-commercials sources such as MetaMorph (Molecular Devices), Imaris (Bitplane), CellProfiler and Bioconductor (8,9) now include modules for automated cell identification. These programs commonly use cellular or nuclear segmentation, i.e., the recognition of connected pixels in binary images obtained from thresholding, to identify individual cells (7,10). However, MNPs have an irregular shape with long dendrites and discontinuous staining on routine histological sections, which makes automated identification of individual cells highly challenging (11). Moreover, only few tools are available that have sought to automate quantitative analysis of cell positioning or that have incorporated tools to assess to what extent observed cell distribution can be considered random.

Here, we have developed a MATLAB-based tissue cytometry application, termed **M**ono**N**uclear **P**hagocyte **m**apping **App**lication (MNPmApp), to perform three important image analysis tasks: (I) Identification of MNPs in mucosal tissue sections based on cell surface staining; (II) measurement of MNP-epithelial and MNP-MNP distances to assess cell-cell interactions; and (III) Monte Carlo modeling to determine randomness or specificity of cellular distribution within the tissue (12,13). Functional interactions between cells within a tissue involve specific molecular mechanisms that are non-random. To quantitatively determine the specificity or randomness of cell interactions on microscopic images, various test statistics such as pairwise inter-cell distances and cell-to-epithelium distances have previously been used. Händel et al. (14) developed an equation that assumes up to six potential contacts between a cell of interest and surrounding cells to predict the expected distribution of two cell types within a tissue. When irregular tissue geometries such as the gastric lamina propria need to be considered, this analytic approach becomes prohibitively complex. Therefore, we included a Monte Carlo modeling feature to compute randomized cell patterns that are compared to the observed cell patterns in our image analysis approach.

Using MNPmApp to analyze a tissue microarray with sections from healthy and *H. pylori-*infected adults, we here demonstrate that, interestingly, observed MNP-epithelial interactions were not significantly different from a randomized distribution, which suggests that non-epithelial cues as well as tissue geometry may control MNP distribution in the human gastric mucosa. However, mean MNP-MNP distances were significantly lower than expected, consistent with MNP cluster formation. Surprisingly, presence or absence of *H. pylori* infection did not significantly alter MNP density or distribution in the tissues. Overall, our analyses demonstrate MNPmApp is a valuable tool for automated, unbiased high throughput analysis of MNP density and distribution in immunofluorescently labeled mucosal tissue sections.

## Materials and Methods

### Tissue samples

Endoscopic gastric biopsies (corpus region) were obtained with local IRB approval from 25 adult and 120 juvenile subjects with abdominal symptoms residing in Santiago, Chile, as previously described (3). Exclusion criteria included (a) use of antibiotics, antacid, H2-blocker, proton-pump inhibitor, bismuth compound, non-steroidal anti-inflammatory drug or immunosuppressive agent during the two weeks prior to endoscopy; and (b) stool examination positive for ova or parasites. *H. pylori* status was determined by rapid urease test and microscopic evaluation, and a study subject was judged colonized with *H. pylori* if one or both tests were positive for the bacteria. Biopsies were formalin-fixed and then paraffin-embedded into a tissue microarray. Only samples from 19 adult patients with well-preserved tissue morphology were considered for our analyses.

### Immunohistochemical staining protocol

To label MNPs and epithelial cells using immunofluorescence, slides were deparaffinized and rehydrated using 5 min washes in differently concentrated ethanol. We then performed heat-induced epitope retrieval in a rice steamer with Dako Target Retrieval Solution (DakoCytomation, Santa Clara, CA) at 98°C for 25 min. Slides were then washed with running distilled water until all unmasking solution had been replaced. Following blocking with a commercial blocking solution, slides were rinsed with PBS-0.05% Tween-20 solution and then incubated with primary antibodies for HLA-DR to label MNPs (Abcam, Cambridge, UK; ab 166777, mouse anti-human IgG2b, clone LN-3) and anti-pan keratin to label epithelial cells (Cell Signaling Technology, Danvers, MA; #4545, mouse anti-human IgG1 clone C11) in a humidity chamber for at least 2 h. After washing slides, secondary antibodies (Southern Biotechnologies, Birmingham, AL) were added for 30 min (HLA-DR: goat anti mouse IgG2b Alexa 488; cytokeratin: goat anti mouse IgG1 biotin followed by streptavidin PE). Cell nuclei were labelled with DAPI (4′,6-diamidino-2-phenylindole). A control slide was processed by labeling with secondary antibodies only to assess background fluorescence. The slides were washed and cover-slipped with Fluoroshield histology mounting medium (Abcam) and sealed with nail varnish prior to microscopic analysis.

### Microscopy

Images were acquired at 20x objective magnification on a Nikon Eclipse T2000-U microscope equipped with a CoolSnap ES digital camera and NIS Elements BR2.30 software (Nikon, Tokyo, Japan). All image analyses were performed using the same set of digital images (n=57) obtained from the 19 adult tissue samples from the tissue microarray described above. Image files were saved as 16 bit *.tif files. Regions of interest (ROIs) consisting of the gastric lamina propria were traced using the selection brush tool in ImageJ (15), version 1.53c and were saved as *.csv files. Borders between the selected lamina propria regions and the epithelium correspond to the epithelial basement membrane that separates lamina propria and epithelium.

### MNPmApp design in MATLAB

The MNPmApp was designed in MATLAB (MathWorks Inc., Natick, MA). MNPmApp uses image processing techniques to identify immunofluorescently labeled MNPs in digital histology images based on the presence of the fluorescent MNP label in the region surrounding a fluorescently labeled cell nucleus. To perform this analysis, first we localized putative nuclear centers by convolving the cell nuclei channel with a pre-tuned Difference of Gaussian (DoG) filter. This template matching filter resulted in a processed image with strong positive responses where stained objects of the expected nuclear size, based on two DoG filters set to a standard deviation of 5 and 6 pixels, were present in the DAPI channel. Next, we applied MATLAB’s “imregionalmax” function to this DoG-filtered image, which identified local maxima, indicating likely centers of nuclei. While this process robustly located the nuclear centers, it also found maxima outside of the ROI and spurious low intensity local maxima due to DAPI background noise. Therefore, the local maxima were further filtered, first using the selected ROI. Second, spurious false positives were removed by filtering the remaining candidate locations using the original cell nuclei channel so that only local maxima with sufficiently strong DAPI signal and template match were retained. Thus, a local maximum of the nuclear detection process was rejected as nucleus location if the DAPI signal at that pixel was below half the median DAPI signal in the image (insufficient signal strength) or if the nuclear detection signal was below 5% of the median DAPI signal in the image (insufficient shape match). The resulting list of maxima was then used as the presumed centers of the nuclei contained inside the ROI.

After the nuclei centers were isolated, the MNPmApp determined which of the nuclei centers were likely associated with MNPs. Association was presumed if an “MNP” signal above the defined threshold level was found in a specified area (defined by the disk size) around a nucleus location. To this end, we integrated the MNP channel around the location of each nuclear center giving a measure of how much MNP signal was near each nucleus. We did this by convolving the MNP channel with a binary disk of user-specified size, which related to the expected spatial extent of one MNP. Finally, using the user’s chosen thresholding parameter, we classified the nuclear centers as MNPs if the disk-convolved MNP signal locally surpassed the threshold. The end result was an x-y coordinate table of the locations of classified MNP nuclei indicating how many MNPs were found within the ROI.

### MNPmApp distance measurements and Monte-Carlo simulations

Based on the pixel coordinates of the identified MNPs, and the ROI delineation of the epithelium, the app then proceeded to compute distances: the Euclidean distance between each MNP and the closest pixel of the epithelial boundary (MNP-EP), as well as the distance between each MNP and its closest peer (MNP-MNP). The app also reports the minimum, maximum and mean for both distances within one image, as well as histograms that show data distribution. For statistical comparison, i.e., to rule out complete spatial randomness, the app then repeatedly creates artificial MNP placements by randomly sampling uniformly (without replacement) an equal amount of points within the ROI, generating 1,000 randomized data sets for each image. MNP-epithelium and MNP-MNP distances are then computed for each of these random datasets. Statistics and histograms of these Monte-Carlo-simulated background distributions are provided as output for further statistical analysis.

### Image processing with MNPmApp

The MNPmApp is available on GitHub (https://github.com/dzosso/Spatial-Stats-App). To avoid a local MATLAB installation, we uploaded the app to a MATLAB online repository for cloud-based computing. For image processing, the file types .tif (microscopy images) and .csv (for ROI data) for each image were input into the MNPmApp. The appropriate image channels were associated with the MNPs (HLA-DR, green) and cell nuclei (DAPI, blue). Threshold and disk size values were adjusted to reflect previously identified optimum values of 0.1 and 6, respectively. The appropriate conversion factor for digital images was used to convert pixels to micrometers. Image processing was then initiated with use of the run feature. A sample input form and a sample data output window are shown in **Supplemental Fig. 1**.

### Manual cell counting

Manual cell counts were performed using the Cell Counter plugin in ImageJ. For optimization and validation of MNPmApp, 3-4 images were selected for generation of receiver operating characteristics (ROC) curves with different threshold and disk size settings, and 12 images were analyzed as the gold standard dataset to determine sensitivity and specificity of MNP identification by MNPmApp. For these analyses, MNPs were identified based on the presence of green fluorescent stain around a central nucleus by JS. In addition, previously obtained cell counts performed on the complete set of 57 images by two other researchers (DB and JD) that were based on identification of green fluorescent signal alone were used for comparison.

### Statistical analysis

Data were analyzed using GraphPad Prism version 9 (GraphPad Software, San Diego, CA, USA) or MATLAB. For each donor, data from 1 – 5 individual digital images covering the entire tissue area available on the tissue array were averaged and expressed as a single data point. Results are presented as individual data points and/or mean ± standard deviation (SD). Differences between values were analyzed for statistical significance by Student’s *t* test, the non-parametric Kruskal-Wallis test, one-way or two-way ANOVA with appropriate multiple comparisons tests. Differences were considered significant at *P*≤0.05. In addition, the Kolmogorov-Smirnov test and the Pearson’s χ^2^ test were used to compare distance distributions between observed and randomized datasets. Z scores were calculated as follows: (observed mean distance - mean of simulated mean distances) / (standard deviation of simulated mean distances). Sensitivity and specificity of MNP recognition by MNPmApp were calculated based on manual counts performed by a researcher (J.S.) used as the gold standard for true positives (TP) and true negatives (TN). Events recognized by MNPmApp, but not the researcher were considered false positives (FP) and events recognized by the researcher, but not by MNPmApp were considered false negatives.

## Results

### Development and validation of an image analysis application (MNPmApp) to assess spatial distribution of MNPs in human mucosal tissue sections

In our previous studies, we showed that MNPs in human gastric mucosa communicate with the gastric epithelium through direct molecular interactions and soluble mediators and that these interactions are impacted by *H. pylori* infection (3,16–18). To analyze the interactions of MNPs with the gastric epithelium as well as with other MNPs *in situ*, we sought to develop an automated digital analysis tool to perform unbiased, statistically relevant high throughput image analyses. As previously described, gastric MNPs were identified based on immunofluorescence labeling for the major histocompatibility class II molecule HLA-DR and were found either directly adjacent to the epithelium (**Fig. 1A,B**) or were distributed throughout the lamina propria (**Fig. 1A,C**). The MNPs frequently formed aggregates or clusters with direct contacts between individual MNPs (**Fig. 1D,E**), whereas other regions contained only solitary MNPs (**Fig. 1F**). The image analysis tool MNPmApp was designed to quantify these different spatial distribution patterns of the MNPs. Since MNPs have an irregular morphology with varying numbers of dendrites, MNP-specific HLA-DR signal intensity was integrated across a concentric disk area with each cell nucleus used as the center, and data points with signals above a defined threshold were identified as MNPs (**Fig. 1G**). Distances between identified MNPs and the manually selected basolateral border of the epithelium and distances between identified MNPs and the nearest other MNP were determined using “nearest neighbor” measurements (**Fig. 1F**).

**Figure 1:**
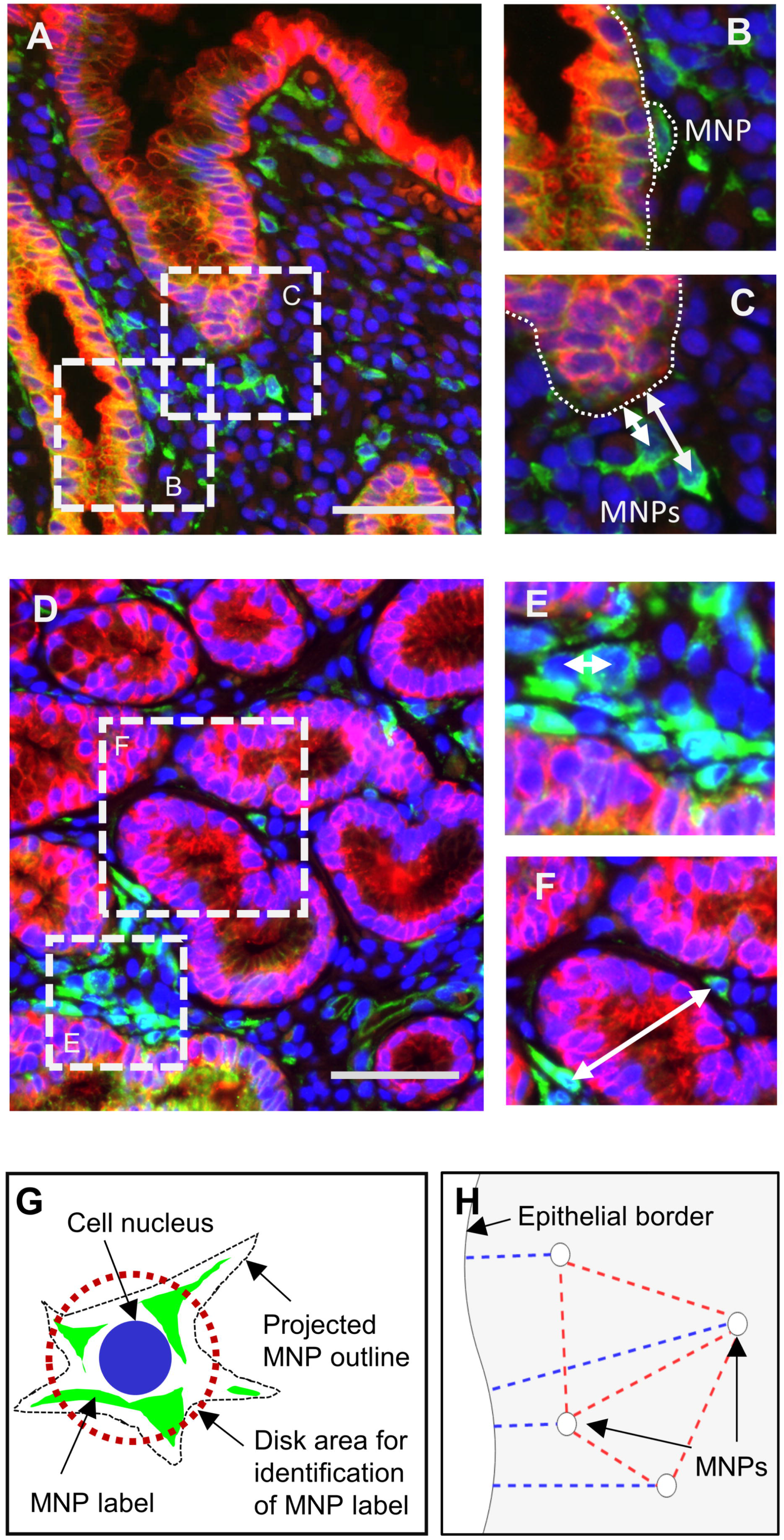
Analysis of MNP positioning within the human gastric mucosa. (**A**) MHC-II-positive MNPs (green) are positioned at variable distances from the gastric epithelium (cytokeratin-positive, red). Cell nuclei are labeled with DAPI (blue). Bar = 50 μm. (**B**) Sample MNP positioned at a larger distance from the epithelium, indicated by the white dashed line. (**C**) Sample MNP positioned directly adjacent to the epithelium. White dashed line indicates position of the basement membrane. (**D**) Some MNPs form cell clusters within the gastric mucosa, with very small distances between individual MNPs. Bar = 50 μm. (**E**) Sample MNP cluster with MNP-MNP distances close to zero. (**F**) Solitary MNP with >50 μm of distance between this MNP and its nearest neighbor. (**G**) MNP identification approach based on position of cell nuclei and surrounding HLA-DR labeling (disk size). Staining intensity for the MNP label is integrated using the selected concentric area around the nuclear center. (**H**) Measurement of MNP-epithelial distances (blue lines) and MNP-MNP distances (red dashed lines) using the “nearest neighbor” approach.

### Processing of digital images with MNPmApp

Digital images of gastric tissues immunofluorescently stained for HLA-DR, cytokeratin, and cell nuclei were prepared for analysis with the MNPmApp by manual selection of the basal epithelial border using the brush selection tool in ImageJ, which created a region of interest (ROI) (**Fig. 2A**). Image processing by MNPmApp then involved splitting a multicolored tif. image into the individual color channels (**Fig. 2B-D**). Next, all cell nuclei present within the selected ROI (**Fig. 2E**) were identified and a yellow marker was placed in the center of each nucleus (**Fig. 2F**). The final step involved the identification of HLA-DR signal that exceeded a certain threshold within a defined area (disk size) around the nuclear centers (**Fig. 2G**). An output image showing the selected epithelial border as well as the identified MNPs, labeled with pink dots, was then generated for verification of MNPmApp performance (**Fig. 2H**).

**Figure 2:**
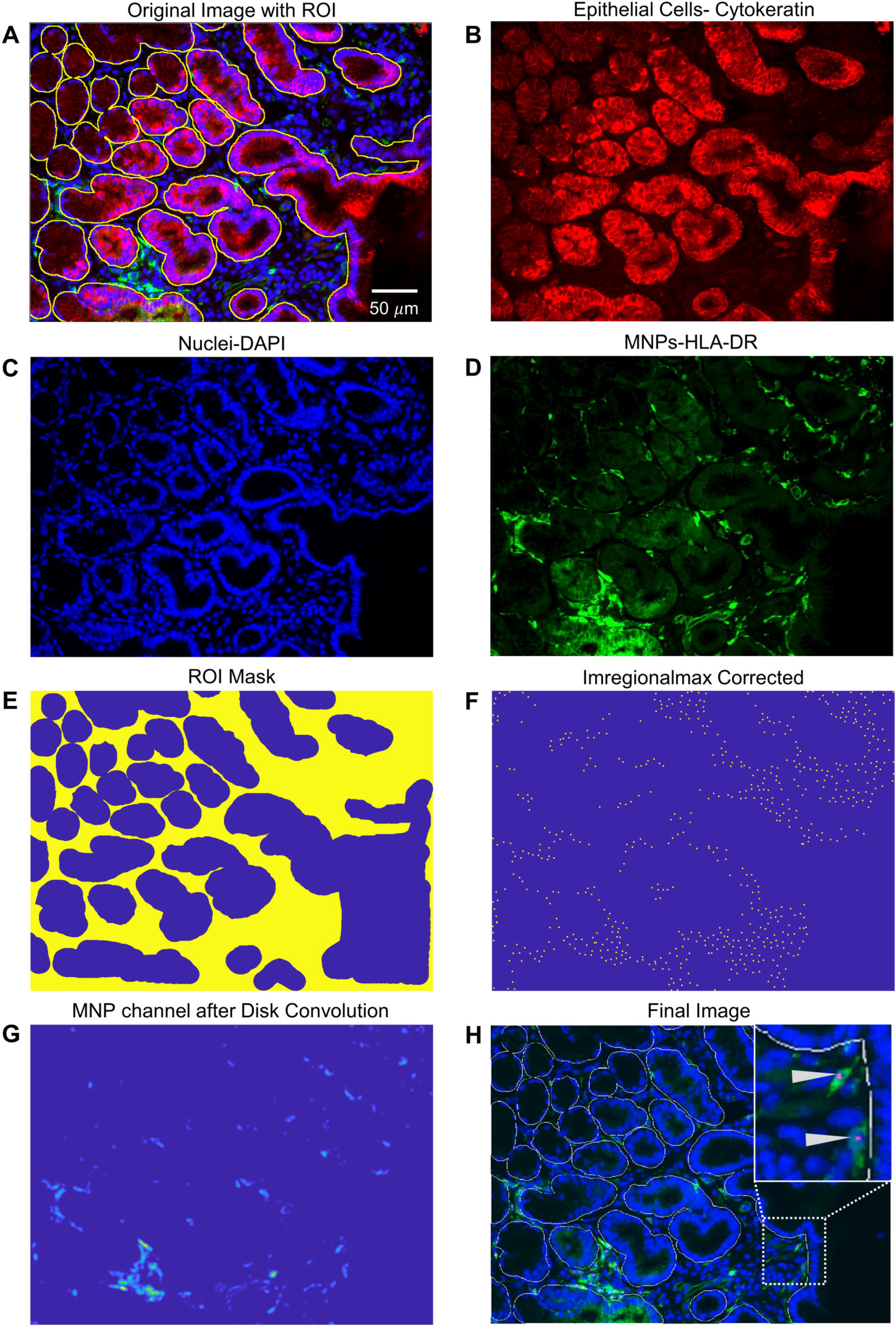
Image processing for automated identification of MNPs using MNPmApp. (**A**) Original merged image showing a paraffin-embedded section of human gastric mucosa with HLA-DR-positive mononuclear phagocytes (green), cytokeratin-positive epithelial cells (red) and cell nuclei (DAPI, blue). Manually generated yellow line indicate the position of the epithelial basement membrane, separating the epithelium from the lamina propria (region of interest, ROI). Bar = 50 μm. (**B**) Single color image showing cytokeratin-positive epithelial cells (red). (**C**) Single color image showing DAPI-positive nuclei (blue). (**D**) Single color image showing HLA-DR-positive MNPs (green). (**E**) Region of interest (ROI) mask outlining the gastric lamina propria in yellow and the epithelium and acellular areas of the slide in blue. (**F**) Position of automatically identified nuclei in the lamina propria. Nuclear centers are indicated by yellow dots. (**G**) Position of automatically identified MNPs. Brighter coloring indicates more intense HLA-DR expression. (**H**) Processed image showing epithelial outlines, blue nuclei and green MNPs. Identified MNPs are labeled with a pink dot.

### Optimization and validation of MNP recognition by MNPmApp

We next sought to optimize the automated MNP recognition by MNPmApp using different settings for threshold and disk size. To that end, three representative images were selected, and MNP identification by MNPmApp was compared to manual identification at 14 different threshold settings (range: 0-0.5; **Fig. 3A,C**). For this gold standard dataset, only MNPs with a strong HLA-DR signal associated with a DAPI-positive nucleus were counted to replicate the criteria used by MNPmApp. Cells identified manually and by MNPmApp also were compared using 10 different disk size settings (range: 1-10) in four individual images (**Fig. 3B,D**). Receiver operating characteristic curves (ROCs) (19) were plotted, and points closest to the top left hand corner of the plots were selected as the best compromise between sensitivity and specificity (red symbols, **Fig. 3C,D**). These points corresponded to a threshold of 0.1 and a disk size of 6. A further comparison between manual counts and automated counts performed by MNPmApp using the predetermined optimum threshold and disk size values on 12 microscopic images revealed a sensitivity of 76.4 ± 15.1% (**Fig. 3E**) and a specificity of 98.3 ± 1.6% (**Fig. 3F**). We did not detect a significant difference in sensitivity and specificity of MNP detection between samples from *H. pylori-*infected and non-infected samples (**Fig. 3E,F**). These data show that, with optimized threshold and disk size settings, MNPmApp successfully identifies MNPs in the gastric tissue sections independent of *H. pylori*-infection status.

**Figure 3:**
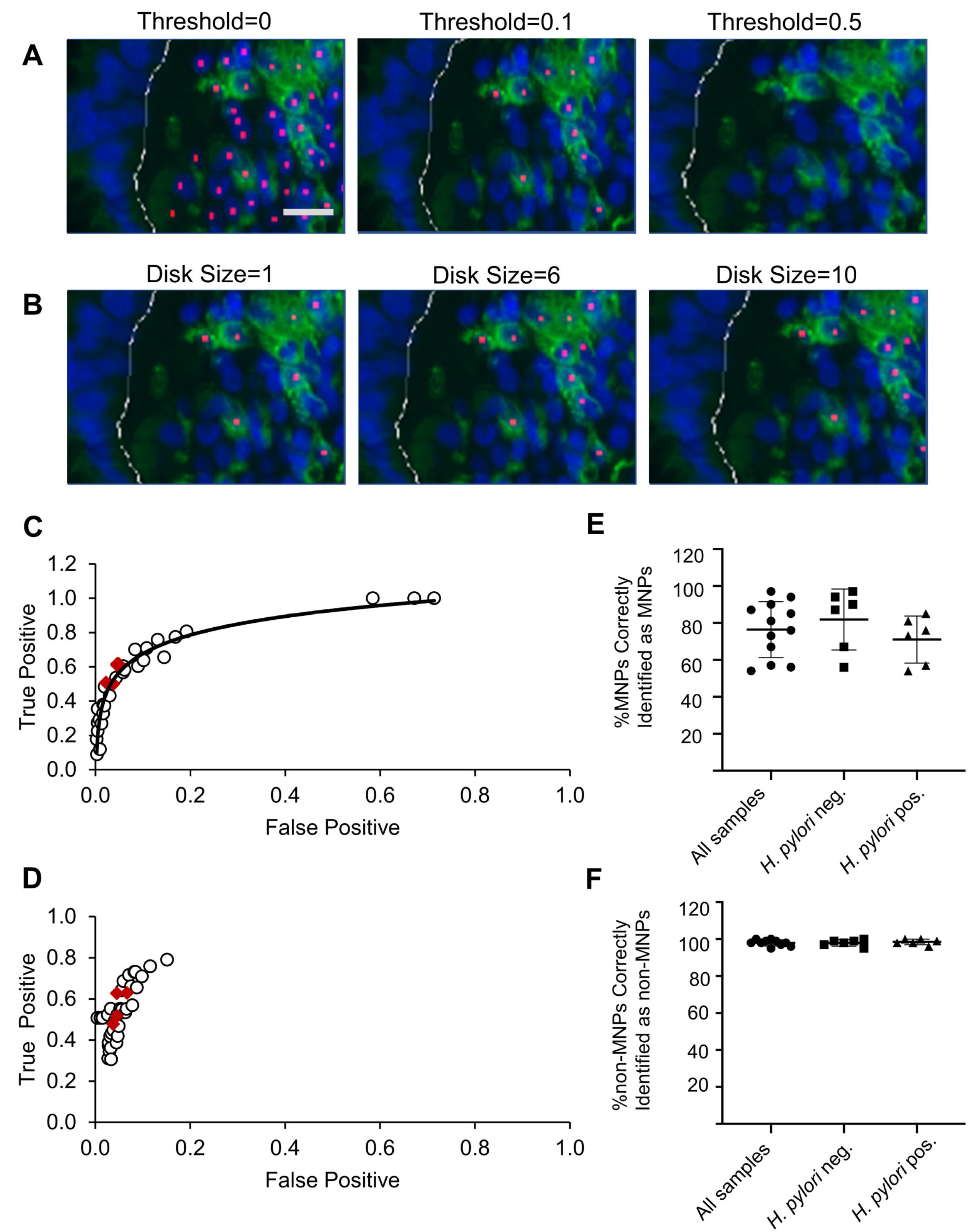
Optimization and validation of MNP identification by MNPmApp. (**A**) Representative images showing MNPs identified by MNPmApp using different threshold values for MNP labeling intensity. Bar = 50 μm. (**B**) Representative images showing MNPs identified by MNPmApp using different disk sizes, which reflects the area of the image around an identified cell nucleus that is analyzed for HLA-DR signal (see Fig 1F). (**C, D**) MNPs identified by MNPmApp were compared to cells manually identified by a researcher (J.S.). True positive and false positive rate for MNPmApp cell identification using (**C**) a range of different threshold values or (**D**) different disk sizes was determined based on manual counting as the gold standard. Dots represent true/false positive rates for three (threshold,C) or four (disk size,D) digital images analyzed with 14 different threshold values and 10 different disk sizes, respectively. Red symbols represent values with optimum MNP identification obtained using a threshold of 0.1 and a disk size of 6. (**E**) Sensitivity and (**F**) specificity of MNP identification by MNPmApp determined by comparing automatically identified to manually identified cells using optimized settings (threshold 0.1, disk size 6). Data were obtained using 12 images of gastric tissues with and without *H. pylori* infection.

### H. pylori infection does not significantly impact MNP density within the gastric mucosa

We next used a data set of 57 digital images from 19 gastric biopsy samples present on the tissue microarray to determine whether *H. pylori* infection alters the number of HLA-DR^+^ MNPs in the human gastric lamina propria. Manual counts previously obtained by two researchers using this dataset were compared to data obtained using MNPmApp. As shown in **Fig. 4A,B**, *H. pylori* infection had no significant impact on the percentage of HLA-DR^+^ MNPs identified by either manual counting or by MNPmApp. Linear regression analysis revealed a significant correlation between manual and automated counts, albeit with a low R^2^ value of 0.29 (**Fig. 4C**). Similar data were obtained when HLA-DR^+^ MNP density in the gastric mucosa was analyzed as cells per tissue area (**Fig. 4D-F**). In general, MNPmApp counted fewer cells than the researchers, likely because cell counts made by the two researchers were based on surface staining alone regardless of whether a nucleus was visible. Interestingly, correlation between data obtained by MNPmApp and manual counts was similar to correlation between manual counts data obtained by two independent researchers (**Fig. 4G,H**). Although we were unable to confirm the *H. pylori*-induced MNP recruitment seen in earlier studies (18,20), our data indicate that image analysis by MNPmApp yields results comparable to those obtained by manual counting.

**Figure 4:**
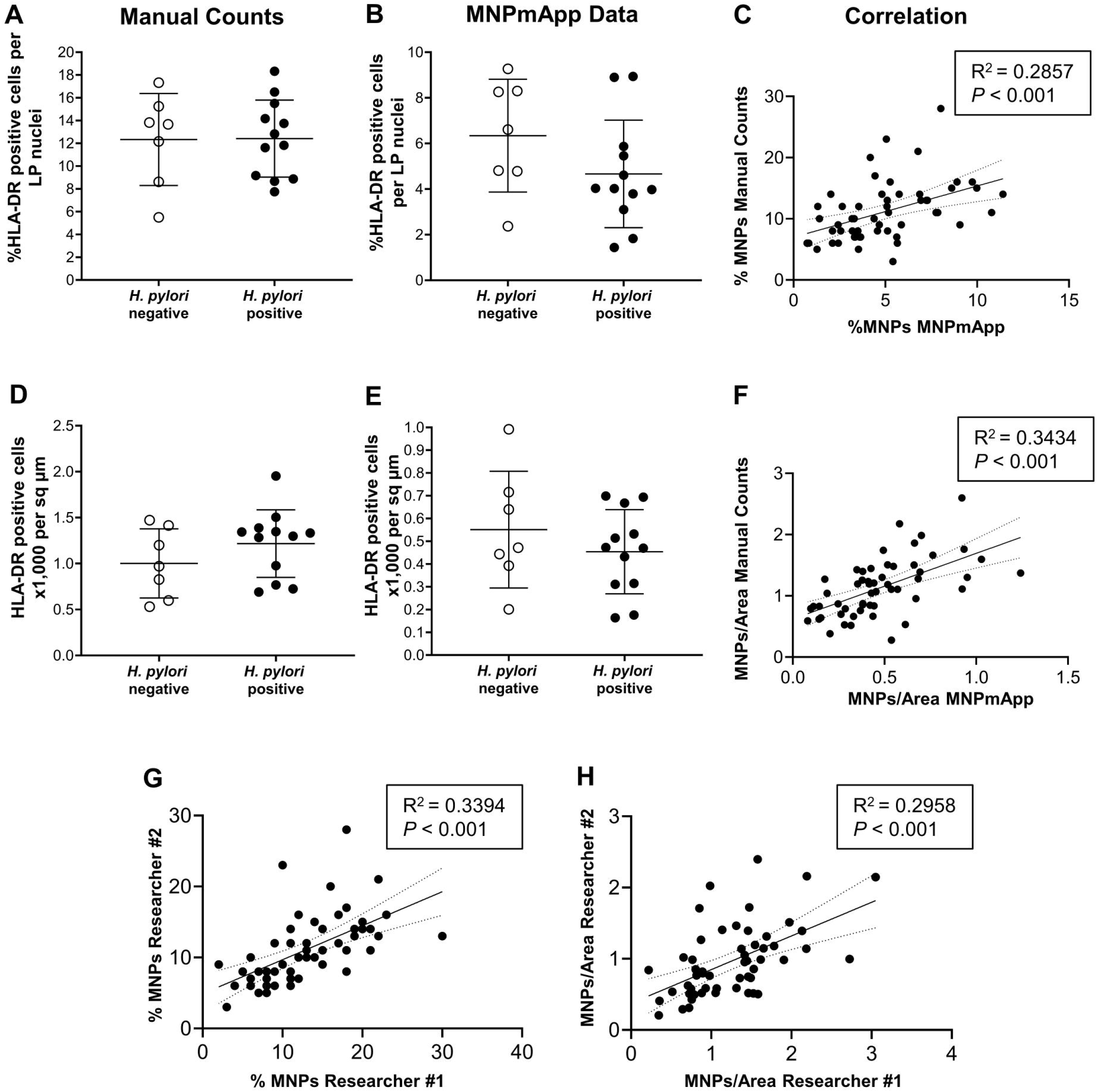
Impact of *H. pylori* infection on MNP density in gastric tissue sections. (**A, B**) Digital images of gastric tissue sections from seven *H. pylori*-negative and twelve *H. pylori*-positive subjects were analyzed for the percentage of HLA-DR^+^ MNPs out of all lamina propria cells using (**A**) manual counting or (**B**) automated identification of MNPs and nuclei with the MNPmApp. Total cell numbers were determined based on total nuclear counts (DAPI channel). Data were analyzed by Student’s *t* test, with no significant difference observed. (**C**) Linear correlation between MNP percentage determined by manual and automated counting; Pearson’s correlation coefficient. (**D, E**) Digital images were analyzed for the number of HLA-DR^+^ MNPs per area of gastric lamina propria using (**D**) manual counting or (**E**) automated identification of MNPs. Tissue area was measured based on the manual selection of the gastric lamina propria (LP) shown in Fig, 2A and E. (**F**) Linear correlation between MNP numbers per area defined as HLA-DR positive cells x1,000 per sq μm based on manual and automated counts; Pearson’s correlation coefficient. (**G**) Linear correlation between MNP percentage determined by two independent researchers using manual counting; Pearson’s correlation coefficient. (**H**) Linear correlation between MNP numbers per tissue area defined as HLA-DR positive cells x1,000 per sq μm determined by two independent researchers using manual counting; Pearson’s correlation coefficient.

### MNPmApp reveals randomized MNP-epithelial distances but smaller than anticipated MNP-MNP distances in human gastric mucosa

In order to quantify MNP distribution within the gastric mucosa, we used MNPmApp to determine the distances between individual MNPs and the epithelium as well as the distances between each MNP and its nearest neighbor. Observed data were compared to 1,000 randomized datasets generated using Monte Carlo modeling for each image. Distance distribution data for a single image are shown in **Fig. 5A, B**. In this particular image, a clear deviation of cellular distance distribution from the simulated distribution pattern was observed. However, mean distances between MNPs and the epithelium for all images did not differ significantly between observed and randomized datasets (**Fig. 5C**). Similarly, a more detailed statistical analysis where we compared observed and simulated MNP-epithelial distance data for individual image files using multiple statistical tests revealed significant differences in only 23-40% of the images (**Supplemental Table 1**). As expected, randomized cell placement resulted in more extreme maximum and minimum MNP-epithelial distances than those that were observed in the tissues. We also observed a strong, highly significant correlation between the observed and randomized data (R^2^ = 0.81, *P<*0.001).

**Figure 5:**
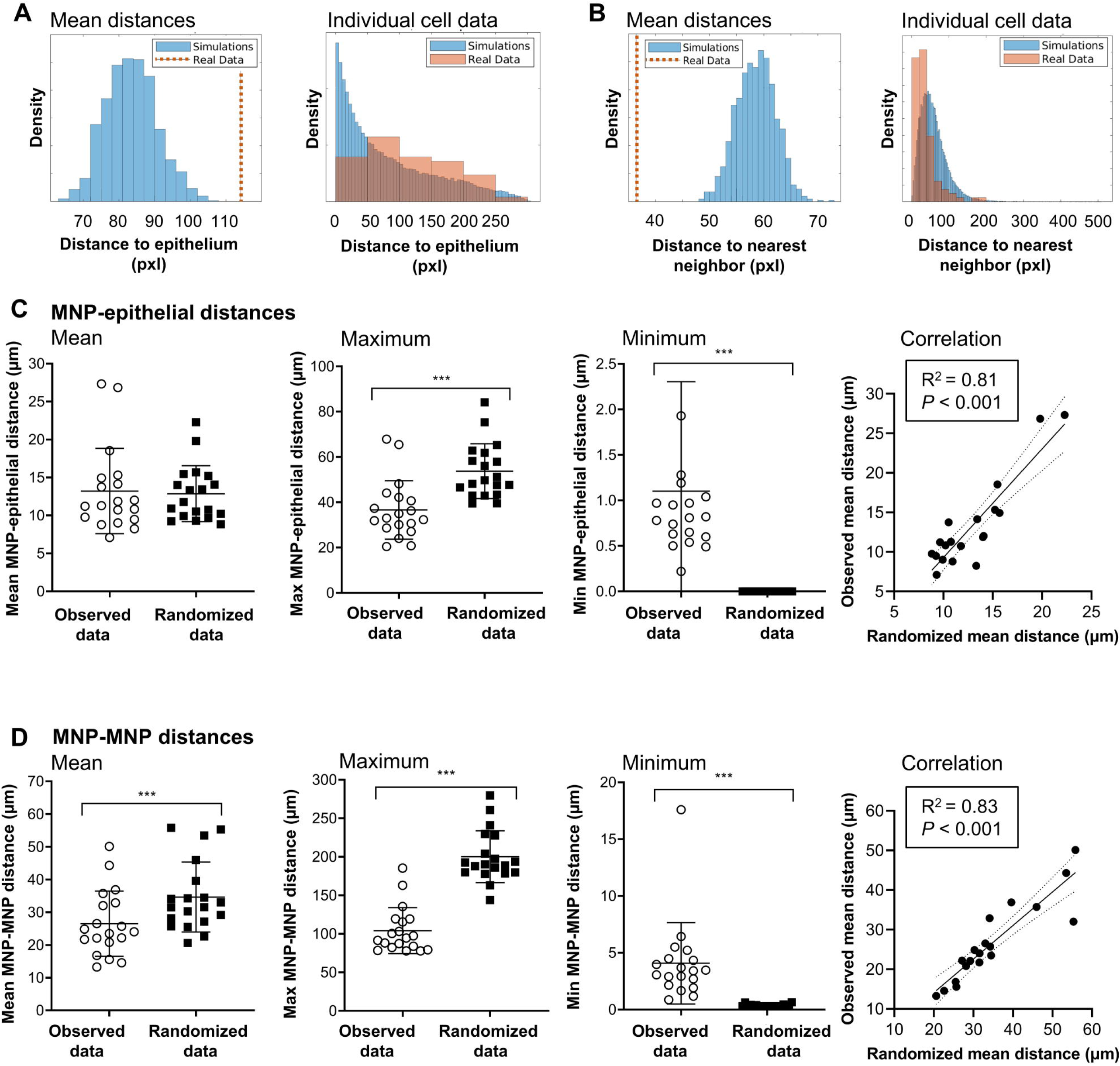
Monte Carlo modeling using MNPmApp reveals random MNP-epithelial distances, but smaller than anticipated MNP-MNP distances. (**A**) Representative distribution plot from one digital image showing the mean observed MNP-epithelial cell distance compared to means of 1,000 randomized sets (left panel) and observed versus randomized MNP-epithelial distances for each individual cell in the image (right panel); (**B**) Representative distribution plot from one digital image showing the mean observed MNP-MNP distance compared to the 1,000 simulated means (left panel) and individual observed versus randomized MNP-MNP distances (right panel); (**C**) Comparison of mean, maximum and minimum MNP-epithelial distances in digital images from n=19 human subjects; mean ± SD. *** Indicates statistically significant differences at *P≤*0.001 (unpaired Student’s *t* test) right panel shows significant correlation between observed and randomized data. (**D**) Comparison of mean, maximum and minimum MNP-MNP distances in digital images from n=19 human subjects; mean ± SD. *** Indicates statistically significant differences at *P≤*0.001 (unpaired Student’s *t* test) right panel shows significant correlation between observed and randomized data.

Interestingly, observed mean distances between MNPs and their nearest neighbors were significantly smaller than distances measured for randomly placed cells (**Fig. 5D**), consistent with the presence of MNP clusters in the tissues, as shown in **Fig. 1D,E**. Significantly lower than expected distances between individual MNPs also were seen for 72-89% of datasets when distance distributions of individual images were analyzed (**Supplemental Table 2**). Again, randomized MNP placement resulted in more extreme minimum and maximum MNP-MNP distances than observed data, and randomized MNP-MNP distances strongly correlated with the observed data (**Fig. 5D**). Overall, the strong correlation and limited differences in cell distribution statistics between real and randomized cell placement within the gastric mucosa indicate that structural features inherent to the tissues likely have the strongest impact on MNP distribution.

To test our original hypothesis that epithelial mediators induced upon *H. pylori* infection recruit MNPs, leading to increased accumulation of MNPs at the epithelial interface in *H. pylori*-positive tissues, we compared actual and randomized MNP distribution in *H. pylori-*infected and non-infected tissues. However, the presence or absence of *H. pylori* infection had no significant impact on either mean MNP-epithelial distances or on MNP-MNP distances, although there was an unexpected trend for increased MNP-epithelial distances in the *H. pylori-*positive samples (**Fig. 6A,B**).

**Figure 6:**
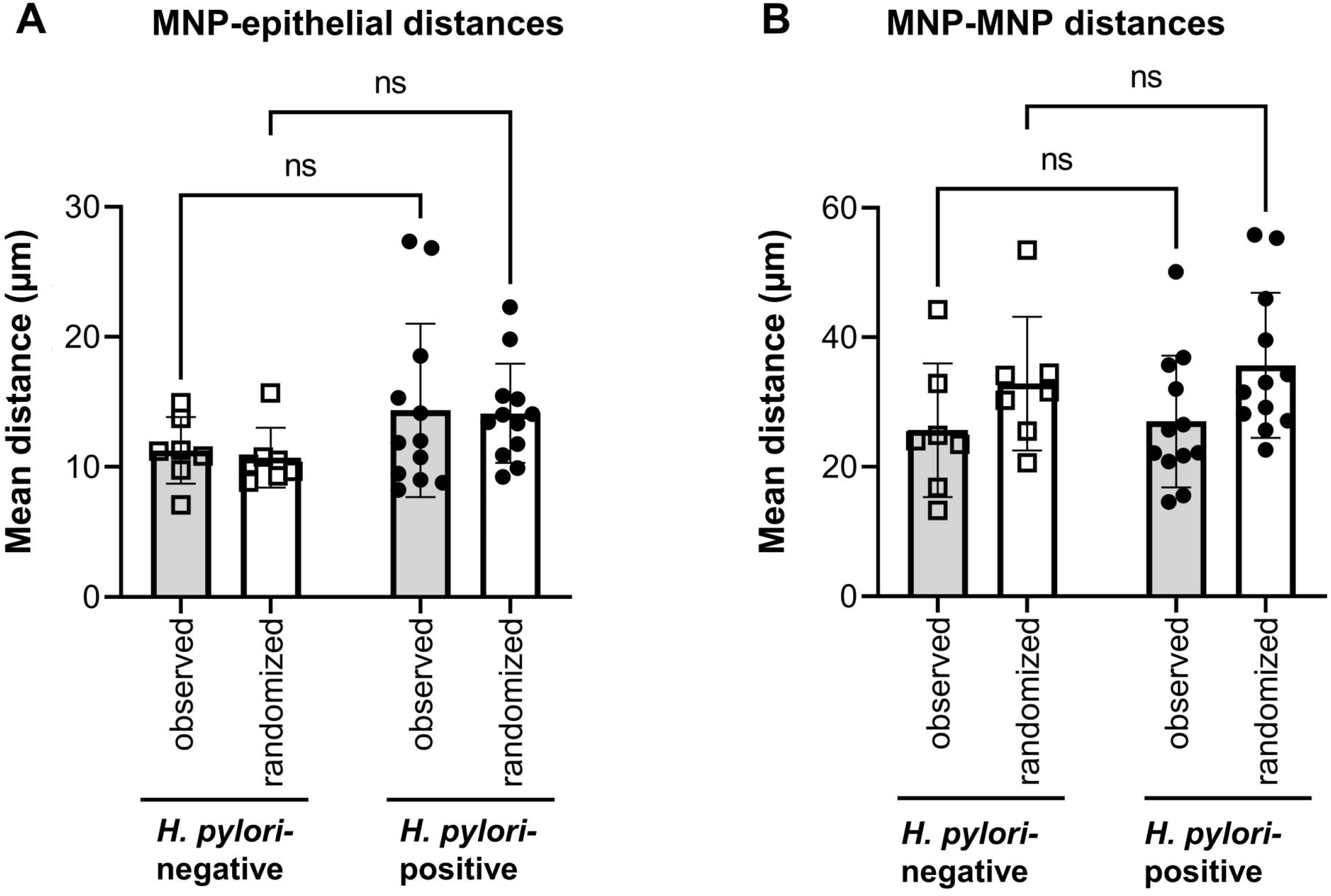
*H. pylori* infection does not significantly alter the distribution of MNPs within the gastric mucosa. Comparison of observed and randomized data generated by Monte Carlo modeling for (**A**) MNP-epithelial distances and (**B**) MNP-MNP distance in healthy (n=7) and *H. pylori-*infected (n=12) human gastric mucosa. Individual data points and mean ± SD are shown; data were analyzed using 2-way ANOVA with Šídák’s multiple comparisons test.

## Discussion

In this study, we have developed and validated an image analysis application, MNPmApp, that maps MNPs in immunohistochemically stained sections of complex tissues. MNPmApp automatically identifies and counts MNPs, determines their spatial relationships with other cells and structures and compares observed MNP distribution with a randomized cell distribution generated by Monte Carlo modeling.

Automated cell identification of cells in immunohistochemical images enables high throughput processing and is less prone to bias than manual analysis performed by investigators (21). However, since MNPs have pleiomorphic shapes with long dendrites, they are difficult to identify using standard digital image analysis algorithms (11). To assess MNP density, we and others have previously used pixel counts, which can accurately identify changes in staining patterns, but do not provide information on cell numbers (20,22). Suberi et al. (23) developed an image processing algorithm to identify and count cells with morphological characteristics of DCs in peripheral blood mononuclear cell samples. DC identification was based on the recognition of characteristic shape signature in phase contrast images of tissue cultures and achieved a sensitivity of 72% and a specificity of 65%. However, this approach is unsuitable for the analysis of tissue sections as dendrites or other membrane extensions may be cropped. Wagner et al. (11) developed an approach to count macrophages in the tumor microenvironment of diffuse large B cell lymphomas based on immunofluorescently labelled tissue sections. The algorithm used a Rudin-Osher-Fatemi (ROF, removes noise from images) filter-based segmentation approach combined with floating intensity thresholding and rule-based feature detection for automated macrophage counting and achieved a >90% correlation with manual counting. In our study, we showed that MNPmApp identifies gastric HLA-DR^+^ MNPs with a high degree of sensitivity and specificity in a gold standard dataset validated by a researcher. Interestingly, we found a highly significant but only moderate correlation between manual and automated count for a larger set of digital images. Notably, correlation between automated analyses and manual counts is frequently poor (24), and correlation between manual counts and automated counts was similar to the correlation between manual counts performed by two independent researchers, indicating that MNP recognition by MNPmApp was adequate, but that our staining protocol may require further optimization to more clearly identify human gastric MNPs.

A key innovative aspect of MNPmApp is that it maps MNP distribution in the gastrointestinal mucosa by measuring distances between identified MNPs and the epithelium and between each MNP and its nearest neighboring MNP. Automated analysis of the spatial distribution of immune cells is an emerging area of research and will ultimately improve our understanding of functional cellular interactions within complex tissues. In a recent study, Tasnim et al. (25) measured the spatial relationship of T cells with cells within lymph nodes using the Pearson correlation coefficient and normalized mutual information, a measure of spatial association that is independent of specific structures but provides some information on randomness. Tasnim’s approach therefore is conceptually similar to our study but lacks a user-friendly application interface. Saylor et al. (10) developed an image analysis algorithm to analyze the organization of myeloid cells and macrophages in the tumor microenvironment and showed a preferential accumulation of CD68^+^ CD163^+^ macrophages in the vicinity of the tumor. Their algorithm was similar to our approach in that it assessed fluorescent signals in donut-shaped areas around cellular nuclei to identify irregularly shaped cells (10). Zwing et al. (26) used a commercially available software (HALO, Indica Labs) to identify multiple immune cell subsets in human colorectal cancer tissues, and distances between cell pairs were determined based on cellular X-Y coordinates, with normalization against myeloid cell density. A comparative study of multiple cell mapping algorithms would be useful to better understand the strengths and weaknesses of the different approaches.

In addition to performing distance measurements, MNPmApp also enabled us to evaluate whether observed cell distributions can be explained by random mechanisms, since observed data were compared to randomized cell placements generated using a Monte Carlo modeling approach (12). Previously, Guidolin et al. (27) successfully used a spatial statistic approach involving computer generation of point patterns that were placed in the area under investigation according to a random (Poisson) distribution (12). We here used a similar approach were each observed cell distribution was compared to 1,000 randomized cell patterns. We hypothesized that mean distances of gastric between gastric MNPs and the epithelial basement membrane would be smaller than mean distances generated using randomized cell placement, based on our previous study that showed chemokine-dependent MNP recruitment by gastric epithelial cells (18). However, our data did not support this hypothesis. Rather, the tight correlation between distance statistics in observed and randomized data sets strongly suggest that structural features of individual tissue sections and ROIs are the major determinants of cellular distribution in morphologically complex tissues. Although MNPs display chemotactic activity towards the gastric epithelium, especially upon *H. pylori* infection, additional mechanisms affecting cell migration and placement in the gastric mucosa also may have contributed to non-significant association between MNPs and the epithelium. For example, MNPs can interact with neuronal or vascular cells present in the mucosa (28), and DCs will routinely leave the lamina propria and migrate through lymphatics towards draining lymph nodes once they have captured antigens (29). Thus, multiple specific interactions between with a variety of cell types likely occur in parallel and may have obscured specific interactions between MNPs and the epithelial layer.

Interestingly, we did observe significantly smaller MNP-MNP distances than predicted based on random cell placement. In the intestine, aggregates of DCs can develop in the lamina propria and attract T cells as a precursor to the development of manifest inflammation in murine transfer colitis (30,31). In the stomach, small aggregates of HLA-DR^+^ MNPs are frequently present, but their functional relevance for gastric homeostasis and disease has not yet been explored. Interestingly, presence or absence of *H. pylori* infection had no significant impact on MNP-MNP distances based on our MNPmApp analyses. Alternative analyses investigating the spatial relationships between MNPs and the vasculature could reveal whether the cells cluster around lymph and blood vessels as pathways for MNP migration into and out of mucosal tissues.

We originally developed the MNPmApp analysis tool to assess whether MNPs in the human stomach are preferentially recruited to the gastric epithelium upon *H. pylori* infection. Contrary to our expectations and previous studies (18,20,32,33), we did not find an increase in MNP density upon *H. pylori* infection in the current dataset, neither using manual counting by two independent researchers, nor using automated cell identification with MNPmApp. There are several possible explanations for these divergent findings: First, since samples were obtained from human subjects with natural infection, disease stage and severity likely vary. Second, distribution of *H. pylori* and the associated inflammatory alterations vary across different regions of the gastric mucosa; therefore, the OLGA approach for diagnostic histopathology scoring recommends the analysis of at least five biopsies from different regions of the stomach (34). Since we used a tissue microarray to minimize differences in staining performance, only one small piece of tissue was included from each donor, which may have excluded tissue regions with more prominent inflammation (35). Lastly, the staining approach we used here was not specific for a particular type of MNP, e.g. DC or macrophages, and may also have labelled B cells. Therefore, changes in the density or spatial distribution of any one specific cell subset may have been masked by other cells identified by the antibody that was used here. Novel labeling techniques enable acquisition of multiplexed fluorescent microscopy images for cell subset identification with more than four markers (36). Future versions of MNPmApp will include additional fluorescence channels, so that cell populations with more complex phenotypes can be identified and distinguished based on marker co-localization.

We had expected smaller MNP-epithelial distances upon *H. pylori* infection, since *H. pylori* infection leads to the induction of epithelial chemokines that attract MNPs (18). Conversely, we saw a trend for increased distances between MNPs and epithelial cells in the *H. pylori-*infected samples. This trend was found in the mean observed MNP-epithelial cell distances seen in *H. pylori* infected versus healthy samples and was also reflected in the differences between the observed compared to the simulated random cell distributions. Possibly, alterations to the gastric lamina propria upon *H. pylori* infection, such as increased recruitment of other inflammatory cells to the epithelial interface (37) or inflammation-induced remodeling of the extracellular matrix (38) may have contributed to the observed changes in average MNP-epithelial distance.

One limitation of the current MNPmApp design was that we did not include an automated approach to identify the epithelium, requiring manual image processing to label the ROIs. Future work will include the development of an optimized staining protocol to delineate the epithelial basement membrane, possibly using laminin staining, so that the epithelium can be automatically identified. One other limitation of our study was that accurate manual counting of cells that formed aggregates was challenging, which made it difficult to determine the sensitivity of automated MNP identification in these areas. Again, including additional fluorescent channels and thus surface markers could help overcome this issue by enabling the identification of smaller cell populations. We have made the current version of MNPmApp available as a shared resource on GitHub, so that it can be utilized by other researchers, and new versions of the app will be deposited in the same location. We anticipate that MNPmApp will be a useful tool for other researchers for quantitative and automated detection of MNPs in histological images.

## Acknowledgements

We gratefully acknowledge funding for this study by NIH grants P30GM110732 (to D. Bimczok and D.Z.), U01EB029242 (to D. Bimczok), P20GM103474 (to D. Bimczok, J.S. and D. Bair) and the M.J. Murdock Charitable Trust Partners in Science Program (to R.A.), and the Simons Foundation collaboration grant for mathematicians #586942 (to D.Z.)

## Author contributions

Conceived and designed the analysis: D. Bimczok, D.Z., C.P.; collected the data: J.S., R.A., J.D.; designed the analysis tools: C.P., D. Bair, J.L., D.Z.; performed the analysis: J.S., D. Bimczok, D.Z.; wrote the original draft of the paper: J.S., D.B., R.A., C.P., reviewed and edited the paper: D. Bimczok, J.S., D. Bair, C.P., J.D., P.R.H., D.Z.; provided clinical samples: P.R.H.

## Figure Legends

**Supplemental Figure 1: MNPmApp user interface**. (**A**) Data input window for file upload and adjustment of threshold, disk size, channel assignments and image resolution. (**B**) Data output screen with summarized data in a *.txt file and single cell data for the observed and randomized cell distances in individual *.csv-files.

**Supplemental Table 1: Image-level data comparing observed versus simulated MNP-epithelial cell distances**. Observed MNP-epithelial cell distance data for each image were compared to randomized datasets from 1,000 simulations using the Kolmogorov-Smirnov, Kruskal-Wallis and Pearson’s χ^2^ test. Table shows individual *P*-values and the Z scores, calculated as (observed mean distance - mean of simulated mean distances) / standard deviation of simulated mean distances. Significant values are labelled green.

**Supplemental Table 2: Image-level data comparing observed versus simulated MNP-MNP distances**. Observed MNP-MNP distance data for each image were compared to randomized datasets from 1,000 simulations using the Kolmogorov-Smirnov, Kruskal-Wallis and Pearson’s χ^2^ test. Table shows individual *P*-values and the Z scores, calculated as (observed mean distance - mean of simulated mean distances) / (standard deviation of simulated mean distances). Significant values are labelled green.

## References

1. Reynolds G, Haniffa M. Human and Mouse Mononuclear Phagocyte Networks: A Tale of Two Species? Front Immunol 2015;6:330.

2. Bimczok D, Grams JM, Stahl RD, Waites KB, Smythies LE, Smith PD. Stromal regulation of human gastric dendritic cells restricts the Th1 response to Helicobacter pylori. Gastroenterology 2011;141:929–38.

3. Bimczok D, Smythies LE, Waites KB, Grams JM, Stahl RD, Mannon PJ, Peter S, Wilcox CM, Harris PR, Das S and others. Helicobacter pylori infection inhibits phagocyte clearance of apoptotic gastric epithelial cells. J Immunol 2013;190:6626–34.

4. Bimczok D, Kao JY, Zhang M, Cochrun S, Mannon P, Peter S, Wilcox CM, Monkemuller KE, Harris PR, Grams JM and others. Human gastric epithelial cells contribute to gastric immune regulation by providing retinoic acid to dendritic cells. Mucosal Immunol 2015;8:533–44.

5. Iliev ID, Spadoni I, Mileti E, Matteoli G, Sonzogni A, Sampietro GM, Foschi D, Caprioli F, Viale G, Rescigno M. Human intestinal epithelial cells promote the differentiation of tolerogenic dendritic cells. Gut 2009;58:1481–9.

6. Smythies LE, Sellers M, Clements RH, Mosteller-Barnum M, Meng G, Benjamin WH, Orenstein JM, Smith PD. Human intestinal macrophages display profound inflammatory anergy despite avid phagocytic and bacteriocidal activity. J. Clin. Invest 2005;115:66–75.

7. Wollmann T, Erfle H, Eils R, Rohr K, Gunkel M. Workflows for microscopy image analysis and cellular phenotyping. J Biotechnol 2017;261:70–75.

8. Smith K, Piccinini F, Balassa T, Koos K, Danka T, Azizpour H, Horvath P. Phenotypic Image Analysis Software Tools for Exploring and Understanding Big Image Data from Cell-Based Assays. Cell Syst 2018;6:636–653.

9. McQuin C, Goodman A, Chernyshev V, Kamentsky L, Cimini BA, Karhohs KW, Doan M, Ding L, Rafelski SM, Thirstrup D and others. CellProfiler 3.0: Next-generation image processing for biology. PLoS Biol 2018;16:e2005970.

10. Saylor J, Ma Z, Goodridge HS, Huang F, Cress AE, Pandol SJ, Shiao SL, Vidal AC, Wu L, Nickols NG and others. Spatial Mapping of Myeloid Cells and Macrophages by Multiplexed Tissue Staining. Front Immunol 2018;9:2925.

11. Wagner M, Hansel R, Reinke S, Richter J, Altenbuchinger M, Braumann UD, Spang R, Loffler M, Klapper W. Automated macrophage counting in DLBCL tissue samples: a ROF filter based approach. Biol Proced Online 2019;21:13.

12. Besag J, Diggle PJ. Simple Monte Carlo Tests for Spatial Pattern. Journal of the Royal Statistical Society. Series C (Applied Statistics) 1977;26:327–333.

13. Diggle PJ, Besag J, Gleaves JT. Statistical-Analysis of Spatial Point Patterns by Means of Distance Methods. Biometrics 1976;32:659–667.

14. Handel N, Brockel A, Heindl M, Klein E, Uhlig HH. Cell-cell-neighborhood relations in tissue sections--a quantitative model for tissue cytometry. Cytometry A 2009;75:356–61.

15. Schneider CA, Rasband WS, Eliceiri KW. NIH Image to ImageJ: 25 years of image analysis. Nat Methods 2012;9:671–5.

16. Roe MM, Swain S, Sebrell TA, Sewell MA, Collins MM, Perrino BA, Smith PD, Smythies LE, Bimczok D. Differential regulation of CD103 (alphaE integrin) expression in human dendritic cells by retinoic acid and Toll-like receptor ligands. J Leukoc Biol 2017;101:1169–1180.

17. Swain S, Roe MM, Sebrell TA, Sidar B, Dankoff J, VanAusdol R, Smythies LE, Smith PD, Bimczok D. CD103 (alphaE Integrin) Undergoes Endosomal Trafficking in Human Dendritic Cells, but Does Not Mediate Epithelial Adhesion. Front Immunol 2018;9:2989.

18. Sebrell TA, Hashimi M, Sidar B, Wilkinson RA, Kirpotina L, Quinn MT, Malkoc Z, Taylor PJ, Wilking JN, Bimczok D. A Novel Gastric Spheroid Co-culture Model Reveals Chemokine-Dependent Recruitment of Human Dendritic Cells to the Gastric Epithelium. Cell Mol Gastroenterol Hepatol 2019;8:157–171 e3.

19. Bray MA, Carpenter A. Advanced Assay Development Guidelines for Image-Based High Content Screening and Analysis. In: Markossian S, Sittampalam GS, Grossman A, Brimacombe K, Arkin M, Auld D, Austin CP, Baell J, Caaveiro JMM, Chung TDY and others, editors. Assay Guidance Manual. Bethesda (MD); 2004.

20. Bimczok D, Clements RH, Waites KB, Novak L, Eckhoff DE, Mannon PJ, Smith PD, Smythies LE. Human primary gastric dendritic cells induce a Th1 response to H. pylori. Mucosal Immunol 2010;3:260–9.

21. Lopez C, Lejeune M, Salvado MT, Escriva P, Bosch R, Pons LE, Alvaro T, Roig J, Cugat X, Baucells J and others. Automated quantification of nuclear immunohistochemical markers with different complexity. Histochem Cell Biol 2008;129:379–87.

22. Inman CF, Singha S, Lewis M, Bradley B, Stokes C, Bailey M. Dendritic cells interact with CD4 T cells in intestinal mucosa. J Leukoc Biol 2010;88:571–8.

23. Suberi AAM, Zakaria WNW, Tomari R. Dendritic Cell Recognition in Computer Aided System for Cancer Immunotherapy. Procedia Computer Science 2017;105:177–182.

24. Inman CF, Rees LE, Barker E, Haverson K, Stokes CR, Bailey M. Validation of computer-assisted, pixel-based analysis of multiple-colour immunofluorescence histology. J Immunol Methods 2005;302:156–67.

25. Tasnim H, Fricke GM, Byrum JR, Sotiris JO, Cannon JL, Moses ME. Quantitative Measurement of Naive T Cell Association With Dendritic Cells, FRCs, and Blood Vessels in Lymph Nodes. Front Immunol 2018;9:1571.

26. Zwing N, Failmezger H, Ooi CH, Hibar DP, Canamero M, Gomes B, Gaire F, Korski K. Analysis of Spatial Organization of Suppressive Myeloid Cells and Effector T Cells in Colorectal Cancer-A Potential Tool for Discovering Prognostic Biomarkers in Clinical Research. Front Immunol 2020;11:550250.

27. Guidolin D, Crivellato E, Nico B, Andreis PG, Nussdorfer GG, Ribatti D. An image analysis of the spatial distribution of perivascular mast cells in human melanoma. Int J Mol Med 2006;17:981–7.

28. Hunyady B, Mezey E, Palkovits M. Gastrointestinal immunology: cell types in the lamina propria--a morphological review. Acta Physiol Hung 2000;87:305–28.

29. Randolph GJ, Angeli V, Swartz MA. Dendritic-cell trafficking to lymph nodes through lymphatic vessels. Nat.Rev.Immunol. 2005;5:617–628.

30. Leithauser F, Meinhardt-Krajina T, Fink K, Wotschke B, Moller P, Reimann J. Foxp3-expressing CD103+ regulatory T cells accumulate in dendritic cell aggregates of the colonic mucosa in murine transfer colitis. Am J Pathol 2006;168:1898–909.

31. Leithauser F, Trobonjaca Z, Moller P, Reimann J. Clustering of colonic lamina propria CD4(+) T cells to subepithelial dendritic cell aggregates precedes the development of colitis in a murine adoptive transfer model. Lab Invest 2001;81:1339–49.

32. Dzierzanowska-Fangrat K, Michalkiewicz J, Cielecka-Kuszyk J, Nowak M, Celinska-Cedro D, Rozynek E, Dzierzanowska D, Crabtree JE. Enhanced gastric IL-18 mRNA expression in Helicobacter pylori-infected children is associated with macrophage infiltration, IL-8, and IL-1 beta mRNA expression. Eur.J Gastroenterol.Hepatol. 2008;20:314–319.

33. Kao JY, Zhang M, Miller MJ, Mills JC, Wang B, Liu M, Eaton KA, Zou W, Berndt BE, Cole TS and others. Helicobacter pylori immune escape is mediated by dendritic cell-induced Treg skewing and Th17 suppression in mice. Gastroenterology 2010;138:1046–54.

34. Rugge M, Pennelli G, Pilozzi E, Fassan M, Ingravallo G, Russo VM, Di Mario F, Gruppo Italiano Patologi Apparato D, Societa Italiana di Anatomia Patologica e Citopatologia Diagnostica/International Academy of Pathology Id. Gastritis: the histology report. Dig Liver Dis 2011;43 Suppl 4:S373-84.

35. Koo M, Squires JM, Ying D, Huang J. Making a Tissue Microarray. Methods Mol Biol 2019;1897:313–323.

36. Blenman KRM, Bosenberg MW. Immune Cell and Cell Cluster Phenotyping, Quantitation, and Visualization Using In Silico Multiplexed Images and Tissue Cytometry. Cytometry Part A 2019;95a:399–410.

37. Blosse A, Lehours P, Wilson KT, Gobert AP. Helicobacter: Inflammation, immunology, and vaccines. Helicobacter 2018;23 Suppl 1:e12517.

38. Sampieri CL. Helicobacter pylori and gastritis: the role of extracellular matrix metalloproteases, their inhibitors, and the disintegrins and metalloproteases--a systematic literature review. Dig Dis Sci 2013;58:2777–83.

