## Supplementary figures and images for "MNPmApp: An image analysis tool to quantify mononuclear phagocyte distribution in mucosal tissues^a, b^"

### Supplemental figure 1

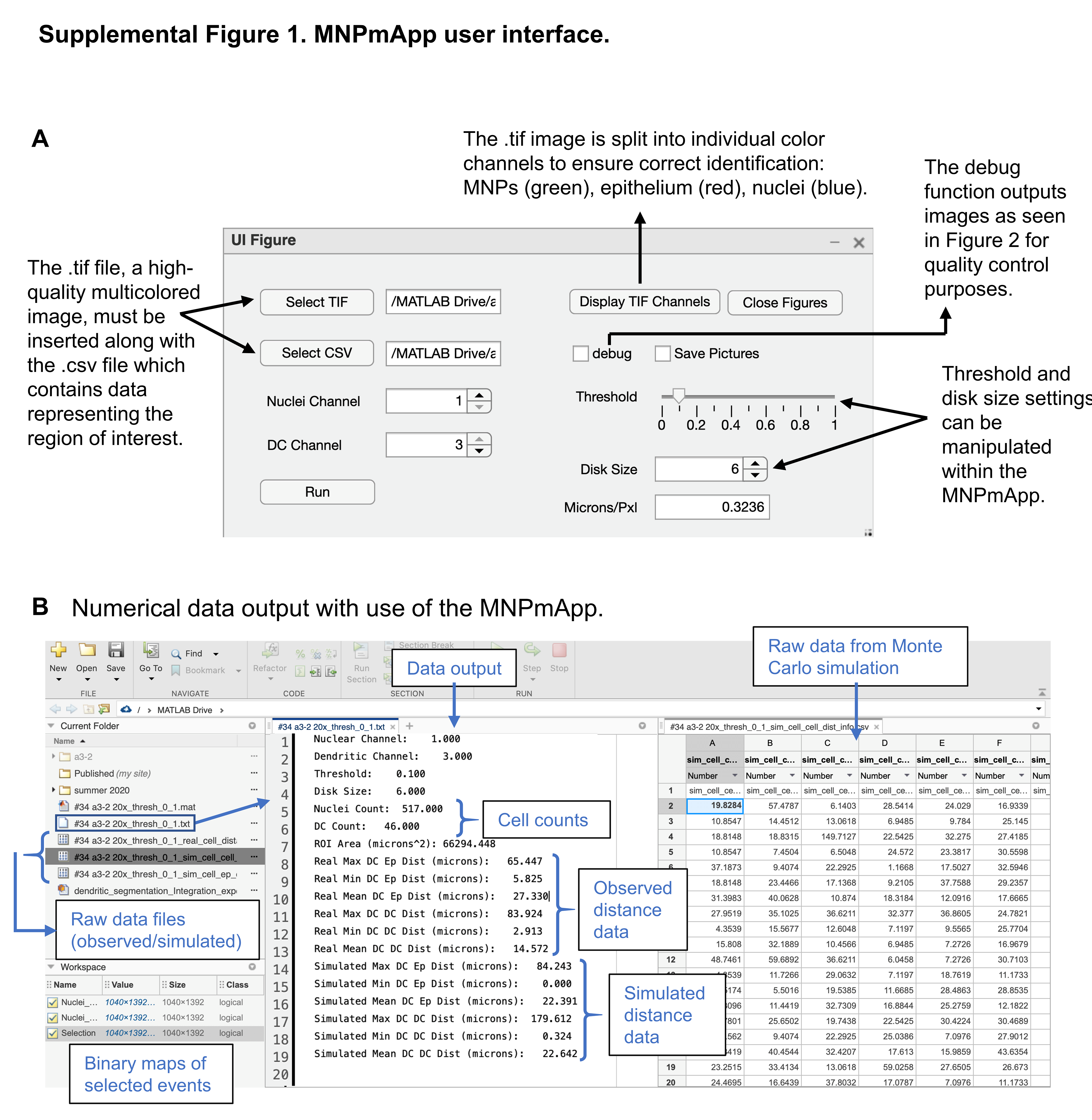
